## Supplementary Information for "Ecological landscapes guide the assembly of optimal microbial communities"

September 21, 2022

**S3 Variance unexplained by higher-order models** **7**

**S4 Simulations of experimental noise** **8**

**S5 Effect of niche overlap on linearity of the map $\vec{N}(\vec{\sigma})$** **8**

**S6 Effect of choice of starting point on search efficacy** **9**

**S7 Supplementary figures** **10**

### 1 Supplementary Information

#### S1 Supplement to Table 1

In this section we provide additional details supplementary to Table 1 in the main text.

**Consumer-resource models with substitutable resources.** Refs. [1–3] show that community assembly under various consumer-resource models with substitutable resources always increase a well-defined mathematical quantity, resembling a Lyapunov function in dynamical systems. This ensures dynamics reach a unique stable steady-state for a given set of starting species, which is the steady-state maximizing the Lyapunov function. Ref. [4] adopts an alternate approach to demonstrate feasibility and local stability of the steady-state.

**Consumer-resource models with cross-feeding or thermodynamic constraints.** In general, there is no mathematical proof guaranteeing a unique stable steady-state for consumer-resource models with cross-feeding or thermodynamic constraints. However, simulations in Refs. [3, 5] find that communities converge to a unique stable steady-state. A key factor in guaranteeing a unique steady-state is possibly the energetic constraints in the models that prevent species from producing resources with more energy than the ones they consumed. Ref. [4] provides some sufficient conditions for feasibility and local stability of consumer-resource models with cross-feeding.

**Many Lotka-Volterra models including those with random interactions with low to moderate variance.** Ref. [6] discusses Lotka-Volterra models that have a Lyapunov function for their dynamics that causes dynamics to converge to a unique steady-state. Refs. [7–9] study Lotka-Volterra models with random interactions using statistical physics techniques. They obtain a phase diagram, which shows that Lotka-Volterra models where interactions have low to moderate variance converge to a unique stable steady-state.

**Empirical observations compatible with a unique stable steady-state.** Refs. [10–12] describe experiments where results are compatible with complex microbial communities converging to a unique stable steady-state. Ref. [13] shows that time-series data from the human gut microbiome is consistent with species having a stable steady-state abundance; abundance fluctuations arise from demographic stochasticity around the steady-state abundance. Ref. [14] shows convergence to a unique stable steady-state occurs when communities are supplied with low nutrient levels, potentially resembling behavior in the theoretical study [7].

**Consumer resource models with non-substitutable resources or species consuming resources diauxically.** Ref. [15] studies consumer-resource models with two non-substitutable resources and finds that multi-stability can occur at certain ratios of resource supply. The maximum fraction of multistable points they obtained, upon fine-tuning resource supply rates, was 15%; the fraction of multistable points dropped rapidly away from this resource supply point. Ref. [16] studies species exhibiting diauxie, i.e., consuming resources in a sequential manner according to their preferences. They show that communities converge to a unique stable steady-state if species have a resource preference order than aligns with the growth rates on each resource. A necessary condition for multistability is the presence of 'anomalous' species i.e., species that consumes resources in an order different from its growth rate on the resources. They then show that anomalous species are evolutionarily disadvantageous and hence unlikely to be found in a community. This makes the chance of multistability very small, albeit difficult to quantify.

**Empirical observations compatible with rare multistability.** Ref. [17] studied communities of 93 different combinations of 8 soil bacterial species. Only 1 of the 93 communitie ($\approx$ 1% ) displayed multistability.

**Lotka-Volterra models with random interactions with large variance or population sub-division.** Ref. [7] study Lotka-Volterra models with random interactions using statistical physics techniques. They obtain a phase diagram, which shows that Lotka-Volterra models where interactions have high variance generically display multiple steady-state attractors with potentially unstable directions. The stochastic invasions and extinctions give rise to complex dynamics, resembling chaos, where the system is switching between the many stable states. Ref. [18] studies stochastic Lotka-Volterra models with anti-symmetric interactions where the populations are divided into many islands. The asynchronous and stochastic dynamics on the many islands ensure species don't go extinct globally and make the total population exhibit complex dynamics resembling chaos.

**Stochastic neutral models.** Ref. [19] summarizes the literature on neutral theory, where species are assumed to be identical. Stochastic fluctuations leads to complex dynamics.

**Consumer resource models with highly nonlinear uptake of more than 3 non-substitutable resources.** Refs. [20, 21] show that consumer-resource models where species grow with more than 3 essential resources (growth follows Liebig's law of the minimum) can exhibit complex dynamics such as heteroclinic cycles and chaos.

**Empirical observations compatible with complex dynamics and chaos.** Ref. [22] studied a complex marine community composed of bacteria, phytoplankton, zooplankton, and detritivores over 2300 days. They observed complex population dynamics resembling chaos. Ref. [23] studied a community in a waste-water treatment plant that was open to constant immigration from the outside and environmental fluctuations. They found the behavior of the open system could be explained by a neutral model.

Please note that this section only approximately condenses a vast and complex literature. The interested reader is encouraged to consult the cited references for a more detailed understanding.

### S2 Ruggedness of community function landscapes

We characterized the structure of community function landscapes using a number of ruggedness measures. We presented the three most useful measures in the main text.

#### S2.1 Variance unexplained by a linear model

We fit ecological landscapes of community function, $\mathcal{F}(\vec{\sigma})$, by the linear model

$$\hat{\mathcal{F}}^{(1)}(\vec{\sigma}) = a^{(0)} + \sum_{i=1}^{S} a_i^{(1)} \sigma_i, \tag{S1}$$

where the fit coefficients $a^{(0)}, a_i^{(1)}$ are obtained by least squares regression, which minimizes $(\hat{\mathcal{F}}^{(1)}(\vec{\sigma}) - \mathcal{F}(\vec{\sigma}))^2$.

The fraction of variance unexplained $U$ is given by

$$U = 1 - \frac{\sum_{\vec{\sigma}}(\hat{\mathcal{F}}^{(1)}(\vec{\sigma}) - \bar{\mathcal{F}})^2}{\sum_{\vec{\sigma}}(\mathcal{F}(\vec{\sigma}) - \bar{\mathcal{F}})^2} \tag{S2}$$

where $\bar{\mathcal{F}}$ is mean function; the numerator and denominator are the explained sum of squares and total sum of squares respectively. Note that the variance explained is equal to the coefficient of determination since the mean of the data and fit are equal in least squares regression. $U$ was estimated from a limited number of data points by calculating $\hat{\mathcal{F}}^{(1)}(\vec{\sigma})$ and $U$ using only the sampled data points.

#### S2.2 Roughness-slope ratio, $r/s$

The roughness-slope ratio, $r/s$, compares the fluctuations in community function to the average trend in the function [24, 25]. The roughness $r$ measured the deviation from the best bit linear model in Eq.S13. It is defined as

$$r = \sqrt{\frac{1}{\|\vec{\sigma}\|}\sum_{\vec{\sigma}}(\mathcal{F}(\vec{\sigma}) - \hat{\mathcal{F}}(\vec{\sigma}))^2}, \tag{S3}$$

where $\sum_{\vec{\sigma}} 1$ is the number of points on the landscape. The slope $s$ is given by

$$s = \frac{1}{S}\sum_{i=1}^{S} |a_i^{(1)}|. \tag{S4}$$

Small $r/s$ values correspond to smooth landscapes—a perfectly linear landscape has $r/s = 0$. Large $r/s$ values correspond to rugged landscapes—$r/s \to \infty$ for $S \to \infty$ for a maximally rugged landscape. Since the maximum value of $r/s$ depends on community size, it is used to compare landscapes of the same size only. When analyzing focal species abundance, $S$ is replaced by $S - 1$ since the effective landscape is smaller Eq. S4.

For estimating $r/s$ from a limited number of sampled data points, $\hat{\mathcal{F}}^{(1)}(\vec{\sigma})$ was computed on the sampled points and the sum in Eq. S3 was restricted to the sampled points. For completely flat landscapes, i.e., if community function was constant, we set $r/s$ to zero.

#### S2.3 Nearest-neighbor correlation and correlation length

We calculated the nearest neighbor correlation $Z_{nn}$ according to the following formula:

$$Z_{nn} = \frac{\frac{1}{\|\vec{\sigma}^{nn}\|}\sum_{\vec{\sigma},\vec{\sigma}^{nn}} \mathcal{F}(\vec{\sigma})\mathcal{F}(\vec{\sigma}^{nn}) - \bar{\mathcal{F}}^2}{\sum_{\vec{\sigma}}\left(\mathcal{F}(\vec{\sigma}) - \bar{\mathcal{F}}\right)^2}. \tag{S5}$$

$\vec{\sigma}^{nn}$ refers to the nearest neighbors of $\vec{\sigma}$ on the landscape, i.e., communities obtained by adding or removing a single species, and $\|\vec{\sigma}^{nn}\|$ is the number of nearest neighbors.

Due to the observed exponential decay of the variance decomposition[1], we can define a landscape correlation length, $\xi$, given by

$$\xi = \frac{-1}{\ln Z_{nn}}. \tag{S6}$$

[1] The nearest neighbor correlation $Z_{nn}$ can also be obtained from the Walsh decomposition [26, 27]

$\xi$ specifies the typical number of species one needs to add/remove to reach a new uncorrelated region of the landscape.

#### S2.4 Fraction of neutral directions

The extrema on the landscape were identified from the relative differences in community function associated with a link between a point and its nearest neighbors:

$$\Delta^B = \frac{\mathcal{F}(\vec{\sigma}^B) - \mathcal{F}(\vec{\sigma}^A)}{\mathcal{F}(\vec{\sigma}^A)} \; \forall B : |\vec{\sigma}^B - \vec{\sigma}^A| = 1. \tag{S7}$$

Many links on the ecological landscapes had zero relative difference, either due to a species not being able to invade or the chosen community function being invariant under invasion of the species. The fraction of the links with zero relative difference was recorded as $F_{\mathsf{neut}}$. In simulations, we considered a relative difference of magnitude less than $10^{-4}$ as zero. In experiments, this threshold would need to be determined from estimates of measurement error or deviation between experimental replicates.

For estimating the fraction of neutral directions and the nearest-neighbor correlation, the landscape was sampled in the same manner. First, we randomly chose $L/2$ points, and then selected an unchosen nearest neighbor for each of the chosen points to make a total of $L$ points. If no unchosen nearest neighbor existed, a random point was chosen instead. $F_{\mathsf{neut}}$ and $Z_{nn}$ were calculated on all sampled nearest neighbors.

#### S2.5 Number of maxima on the landscape

We also computed the number of maxima on the landscape, defined as points with community function greater than or equal to its neighbors (strictly greater than at least one neighbor). We used Eq. S7 with a threshold of $10^{-4}$ to compare the community function of neighboring points. Most maxima had neutral links associated with it. And so we also counted the number of unique maxima, defined as maxima where all species initially present survived in the final community. The number of unique maxima on the landscape did not correlate significantly with search success.

#### S2.6 Fitness Distance Correlation, FDC and ranked Fitness Distance Correlation $r_{\mathsf{FDC}}$

The fitness distance correlation, FDC, measures the deviation of the observed landscape from a single-peaked landscape with community function decreasing linearly with distance from the peak [28, 29]. It is given by the Pearson correlation of the community function and hamming distance from the global optimum i.e.,

$$\mathsf{FDC} = \frac{\sum_{\vec{\sigma}} \left(\mathcal{F}(\vec{\sigma}) - \bar{\mathcal{F}}\right)\left(D_{\mathsf{opt}}(\vec{\sigma}) - \bar{D}_{\mathsf{opt}}\right)}{\sqrt{\sum_{\vec{\sigma}} \left(\mathcal{F}(\vec{\sigma}) - \bar{\mathcal{F}}\right)^2}\sqrt{\sum_{\vec{\sigma}} \left(D_{\mathsf{opt}}(\vec{\sigma}) - \bar{D}_{\mathsf{opt}}\right)^2}}, \tag{S8}$$

where $D_{\mathsf{opt}}(\vec{\sigma})$ is the hamming distance to the optimum, $\bar{\mathcal{F}}$ is the average community function, and $\bar{D}_{\mathsf{opt}}$ is the mean distance to optimum.

We also calculated the Ranked Fitness Distance Correlation $r_{\mathsf{FDC}}$, which computed the correlation of fitness ranks to distance. Both FDC and $r_{\mathsf{FDC}}$ vary from $-1$ to 1, with $\mathsf{FDC} = -1$ $(r_{\mathsf{FDC}} = -1)$

corresponding to a linear (nonlinear but monotonic in distance) landscape with a single peak. Both FDC and $r_{\mathsf{FDC}}$ are 0 for an uncorrelated rugged landscape.

We found that these measures tended to be informative only for landscapes of community functions like diversity, which received contributions from all species in the community, and not community functions like the abundance of a focal species.

##### S2.7 Auto-correlation and correlation length

Instead of performing a gradient-ascent search, one can also explore the community function landscape by taking random steps to nearest neighbors.The random walk auto-correlation, $R_{\mathsf{rw}}$, and the landscape auto-correlation function, $R_{\mathsf{l}}$ are defined with respect to this random exploration of the landscape [26, 27]. On an statistically isotropic landscape, the random-walk auto-correlation is given by Pearson correlation between points on a random walk trajectory $s$ steps apart i.e,

$$R_{\mathsf{rw}}(s) = \frac{\langle \mathcal{F}_t \mathcal{F}_{t+s} \rangle - \langle \mathcal{F} \rangle^2}{\langle \mathcal{F}^2 \rangle - \langle \mathcal{F} \rangle^2}, \tag{S9}$$

where $\mathcal{F}_t$ is the community function at step $t$ and $\langle\rangle$ denotes averaging over $t$ over a sufficiently long walk (assuming ergodicity) or averaging over sufficient number of walks with different origins. The landscape autocorrelation is defined as

$$R_{\mathsf{L}}(d) = \frac{\langle \mathcal{F}(\vec{\sigma}_A)\mathcal{F}(\vec{\sigma}_B) \rangle - \langle \mathcal{F} \rangle^2}{\langle \mathcal{F}^2 \rangle - \langle \mathcal{F} \rangle^2} \ \mathsf{s.t.} D(\vec{\sigma}_A, \vec{\sigma}_B) = d, \tag{S10}$$

where $D$ measures the Hamming distance between the two points. The two correlation measures are related by

$$R_{\mathsf{rw}}(s) = \sum_d \phi_{sd} R_{\mathsf{L}}(d), \tag{S11}$$

where $\phi_{sd}$ is the probability that a random walk $s$ steps long is at a distance $d$ away.

For exponentially decaying auto-correlation, we can identify an associated correlation length $l$ defined by the form $R_{\mathsf{rw}}(s) \sim e^{-\frac{s}{l}}$ , and estimated as

$$l = \frac{-1}{\ln\left(R_{\mathsf{rw}}(1)\right)} \tag{S12}$$

##### S2.8 Species-wise Optimizability, SWO

Species-wise Optimizability SWO quantifies the performance of searching through the landscape by choosing the presence/absence of each species independently [29]. We calculated this by:
1. Choose a starting point $A$, with presence-absence vector $\vec{\sigma}^A$.
2. For each species $i$, compare the community function at $A$ with the function obtained by flipping $\sigma_i$. Record all $i$ where flipping $\sigma_i$ increased the community function.
3. Construct the vector $\vec{\sigma}^{A'}$, corresponding to the vector obtained by flipping all recorded $\sigma_i$. 4. Repeat steps $1-3$ for all starting points on the landscape to get sets $\{A\}$ and $\{A'\}$ .

Following this, the Species-wise Optimizability SWO is calculated as the ratio of average pairwise distances between the points in the set $\{A'\}$ and points in the set $\{A\}$. We do not estimate SWO from limited data.

#### S3 Variance unexplained by higher-order models

We fit ecological landscapes of community function $\mathcal{F}(\vec{\sigma})$ incorporating higher-order species interactions using the following class of models:

$$\hat{F}^{(k)}(\vec{\sigma}) = a^{(0)} + \sum_{i=1}^{S} a_i^{(1)}\sigma_i, + \sum_{i=1}^{S}\sum_{j=1}^{i-1} a_{ij}^{(2)}\sigma_i\sigma_j + \dots \mathcal{O}\left(a^{(k)}\right) + \epsilon\left(\vec{\sigma}\right), \tag{S13}$$

where the fit coefficients $a^{(0)}, a^{(1)}, ...$ defined up to order $k$ are obtained by least squares regression minimizing $(\hat{\mathcal{F}}^{(k)}(\vec{\sigma}) - \mathcal{F}(\vec{\sigma}))^2$. (Note that these models are linear from the point of view of regression since all the interaction terms are provided explicitly.)

The fraction of variance unexplained by the model of order $k$ , $U^{(k)}$, is given by

$$U^{(k)} = 1 - \frac{\sum_{\vec{\sigma}}(\hat{\mathcal{F}}^{(k)}(\vec{\sigma}) - \bar{\mathcal{F}})^2}{\sum_{\vec{\sigma}}(\mathcal{F}(\vec{\sigma}) - \bar{\mathcal{F}})^2} \tag{S14}$$

where $\bar{\mathcal{F}}$ is mean function; the numerator and denominator are the explained sum of squares and total sum of squares respectively. The variance explained is equal to the coefficient of determination since the mean of the data and fit are equal in least squares regression.

Since inverting matrices becomes computationally expensive for large landscapes, we used an alternative method to calculate the the fraction of variance unexplained. Transforming from presence-absence basis, $\sigma_i \in \{0, 1\}$, to the orthogonal basis $s_i \in \{-1, 1\}$, allows one to use a technique resembling Fourier transforms, called Walsh decomposition [30, 31]. The Walsh/Fourier decomposition of the community function on the transformed landscape ($s_i \in \{-1, 1\}$ ) is

$$\tilde{\mathcal{F}}(\vec{s}) = \tilde{a}^{(0)} + \sum_{i=1}^{S} \tilde{a}_i^{(1)} s_i, + \sum_{i=1}^{S}\sum_{j=1}^{i-1} \tilde{a}_{ij}^{(2)} s_i s_j + \dots, \tag{S15}$$

where $\tilde{a}$ are the Fourier components and the expansion continues till order $S$. Orthogonality allows one to compute each Fourier component without matrix inversion, for e.g.,

$$\tilde{a}^{(0)} = 2^{-S} \sum_{\vec{s}} \mathcal{F}(\vec{s}), \tag{S16}$$

$$\tilde{a}_i^{(1)} = 2^{-S} \sum_{\vec{s}} s_i \mathcal{F}(\vec{s}), \tag{S17}$$

etc. One can then perform a variance decomposition akin to calculating the power spectrum. This variance decomposition gives us

$$2^{-S}(\tilde{\mathcal{F}} - \bar{\mathcal{F}})^2 = \sum_{i=1}^{S}(\tilde{a}_i^{(1)})^2, + \sum_{i=1}^{S}\sum_{j=1}^{i-1}(\tilde{a}_{ij}^{(2)})^2 + \dots \mathcal{O}\left(a^{(S)}\right) \tag{S18}$$

We can calculate the fraction of the variance explained by a model of order $k$ by truncating the right hand side at order $k$ and normalizing by the total variance i.e.,

$$U^{(k)} = 1 - \frac{1}{\sum_{\vec{\sigma}}(\mathcal{F}(\vec{\sigma}) - \bar{\mathcal{F}})^2}\left[\sum_{i=1}^{S}(\tilde{a}_i^{(1)})^2 + \sum_{i=1}^{S}\sum_{j=1}^{i-1}(\tilde{a}_{ij}^{(2)})^2 + \dots \mathcal{O}\left(a^{(k)}\right)\right] \tag{S19}$$

We imposed a threshold of $10^{-6}$ in the fraction of variance unexplained when plotting and fitting data in Fig 6.

### S4 Simulations of experimental noise

Two forms of experimental noise was simulated to test robustness of ruggedness estimates in the presence of noise. For simulating additive measurement noise, we modified the community functions at each point on the landscape as:

$$\mathcal{F}_M(\vec{\sigma}) = \mathcal{F}(\vec{\sigma}) + \overline{\mathcal{F}}\lambda_M\, \eta(\vec{\sigma}), \tag{S20}$$

where $\lambda_M$ measures the strength of the noise, $\eta$ is a normally distributed random variable, and $\overline{\mathcal{F}}$ is the average community function on the landscape. For simulating multiplicative noise, we modified the community function at each point on the landscape as:

$$\mathcal{F}_X(\vec{\sigma}) = \mathcal{F}(\vec{\sigma}) + \mathcal{F}(\vec{\sigma})\,\lambda_X\, \eta(\vec{\sigma}), \tag{S21}$$

where $\lambda_X$ measures the strength of the noise and $\eta$ is a normally distributed random variable.

### S5 Effect of niche overlap on linearity of the map $\vec{N}(\vec{\sigma})$

The aim of this section is to provide additional intuition, through analytical investigation of simple models, about why the map from species presence-absence to steady-state abundance, $\vec{N}(\vec{\sigma})$, becomes less linear with increasing niche overlap. We begin with a simple consumer-resource model in the limiting scenario of exclusively specialist species, each having a private resource. Without loss of generality, we will assign species $i$ the private resource $i$. At steady state, we have the following equations being satisfied (if the species survives),

$$(C_{ii}R_i^*) - m_i = 0, \tag{S22}$$

$$\tau^{-1}\left(R_i^0 - R_i^*\right) - C_{ii}R_i^*N_i^* = 0, \tag{S23}$$

where $*$ denotes steady-state. Solving this system of equations, we get the steady-state abundances of species to be

$$N_i^* = \frac{R_i^0}{\tau m_i} - \frac{1}{\tau C_{ii}}, \tag{S24}$$

when positive. (If negative the species goes extinct and abundance is zero.) Clearly, species abundance in this limit is independent of all other species present. Hence a linear map can fit steady-state abundances perfectly in the limit of specialist species with no niche overlap.

To see why the linearity of $\vec{N}(\vec{\sigma})$ reduces as number of inter-species interactions increases, we will analyze a generalized Lotka-Volterra model (gLV) in a simple scenario. We choose a gLV model instead of a consumer-resource model for analytical convenience. We will assume that the species interactions (summarized in the interaction matrix $B$) are such that all species present in the starting community survive at steady-state, in all possible starting species combinations from the species pool (indexed by the species presence-absence vector $\vec{\sigma}$). The gLV model equation we consider will be

$$\frac{dN_i}{dt} = N_i \left[ r_i + \sum_j B_{ij} N_j \right]. \tag{S25}$$

The sum over $j$ runs only over species present in the community. Another way of interpreting this equation is that the relevant interaction matrix changes depending on the species present in the community,. In other words, $\vec{\sigma}$ identifies the appropriate sub-matrix of the full interaction matrix, obtained by deleting the rows and columns corresponding to species absent in the community. We will refer to this submatrix as ${}^{\vec{\sigma}}B$.

Without loss of generality, we will focus on a single species, with index 1. For a given initial presence absence vector, the steady-state abundance of species 1, $N_1^*$, will be

$$N_1^*(\vec{\sigma}) = -\sum_j {}^{\vec{\sigma}}B_{1j}^{-1} r_j. \tag{S26}$$

This equation will give us the steady-state abundance $N_1^*$ for any species combinations where species 1 is present initially, such as., $\vec{\sigma} = \{[1,0,0,...],[1,1,0,...],[1,0,1,...],[1,1,1,...]\}$. Similar equations will hold for other species as well.

Given the steady-state abundances, we can now consider how well the steady-state abundances will be fit by the linear model (Eq.1 in the main text). For steady-state abundances in Eq. (S26) to agree with the prediction of the linear model, we need ${}^{\vec{\sigma}}B^{-1}$, when $\vec{\sigma}$ changes, to behave like the single matrix in the linear model $A$. In the zero niche overlap scenario, where the species interacts with only itself, $B$ is diagonal. The inversion of all possible sub-matrices is trivial and the abundance of species will be independent of the presence-absence of other species; the abundance of species 1 will be $N_1^* = -\frac{r_1}{B_{11}}$). Thus the linear model will fit the zero niche overlap scenario perfectly, as in the case of a consumer-resource model of specialists with no niche overlap.

When niche overlap increases, more matrix elements in $B$ are non-zero. The inverses of the different submatrices, ${}^{\vec{\sigma}}B^{-1}$ have a complex, nonlinear relationship in general. For example, the inverse of two matrices describing communities differing by a single species are related by the nonlinear, Woodbury identity [32]. Hence the linear model fits the data poorly.

### S6 Effect of choice of starting point on search efficacy

In the main text, we present the search outcome averaged over all choices of the starting point. We believe this reflects the experimental scenario where information regarding the ideal choice of starting point is limited or even misleading. To evaluate the effect of the starting point on search, we recorded the efficacy of search optimizing Shannon diversity when started from three special choices of the starting community: a community with all species present, a randomly chosen single species present, and a randomly chosen half of the candidate species present. Although intuitively, one expects a higher diversity starting from a community with all species present, the increase in search outcome (over the average search outcome) was very small (0.01, 0.00, and 0.00 for all species, single species, and half species communities respectively, see S12 Fig). Furthermore, the correlation between search outcome and ruggedness remained robust for all three choices of

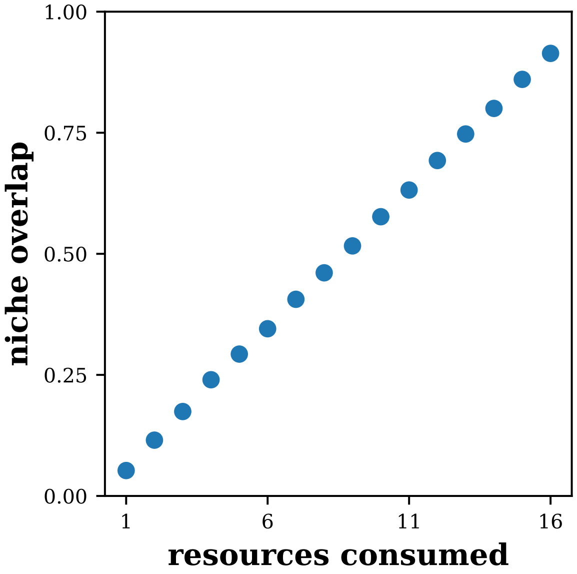

Figure S1: **Niche overlap between species increases with number of resources consumed.** The niche overlap in the 16 species pools shown in Fig 2. Niche overlap was quantified by the average cosine similarity between the consumption vectors of each species pair in the pool, as in previous studies [33]. In these simulations, each non-zero consumption matrix element was drawn from a gamma distribution. The niche overlap increased with number of resources when consumption matrix elements were drawn from other distributions as well (see S11 Fig).

starting communities. Thus, ruggedness is informative of search efficacy from specially chosen starting communities as well.

With intimate knowledge of the system, one can indeed choose special starting points that systematically improve or diminish depending on the community function (at least for simple models). For e.g., consumer-resource models without cross-feeding incorporate only inter-species competition. Therefore, the abundance of a focal species is maximum when it is alone. We are guaranteed to find the optimal community if we start from a community with focal species alone (the optimal community) or one of its nearest-neighbs. It needs to be stressed, however, that this is possible only for specific models and choices of community function—it is unlikely to be relevant for experimental communities.

### S7 Supplementary figures

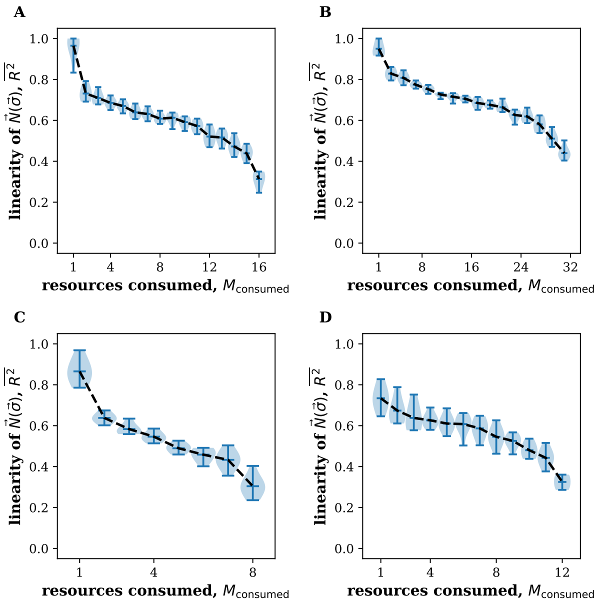

Figure S2: $\vec{N}(\vec{\sigma})$ **is linear for specialists and nonlinear for generalists—independent of the number of resources and presence of cross-feeding.** The map from species presence to steady-state abundances $\vec{N}(\vec{\sigma})$ becomes more nonlinear with increasing niche overlap across a range of models and parameters. The linearity of the map is quantified by $\overline{R^2}$. Niche overlap increases when species in the pool consume more resources. We plot consumer resource models with $S = 16, M_{tot} = 16$ in **(A)** , $S = 16, M_{tot} = 32$ in **(B)** , and $S = 16, M_{tot} = 8$ in **(C)** ; and cross-feeding model with $S = 12, M_{tot} = 12$ in **(D)**. There were 10 replicates for each data point. Simulation parameters are described in Methods.

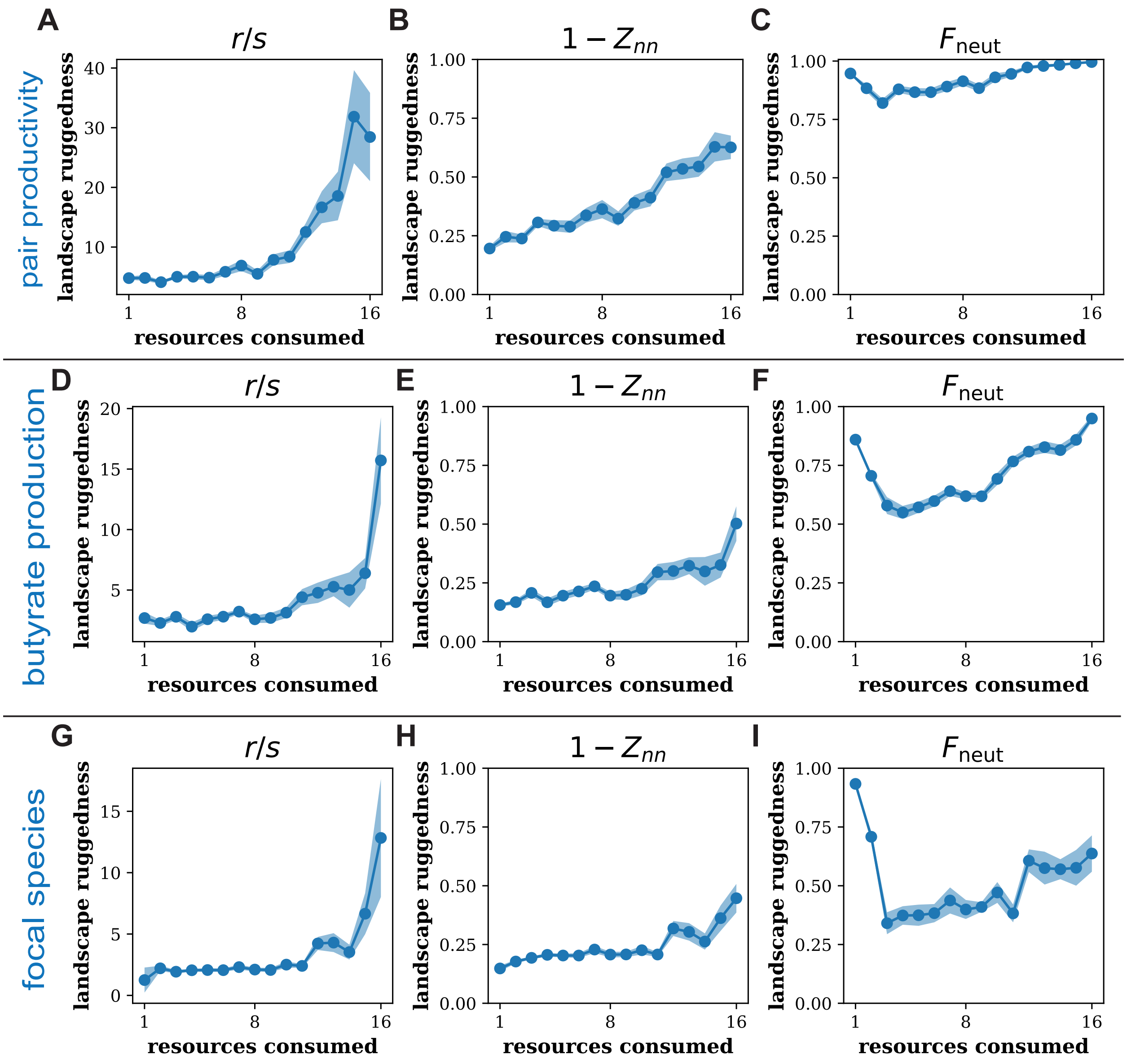

Figure S3: **The ruggedness of ecological landscapes of other community functions.** Panels show $r/s$, $1 - Z_{nn}$, and $F_{\text{neut}}$ the landscapes with community function being pair productivity, butyrate production, and focal species abundance. The fraction of neutral directions $F_{\text{neut}}$ was high at low niche overlap, when all of the species occupied separate niches, because the species (or species pair) responsible for the community function was unaffected by the addition or removal of other species. This causes the community function to be left unchanged and a concordant increase in the number of neutral directions. Simulations were the same as in Fig 5.

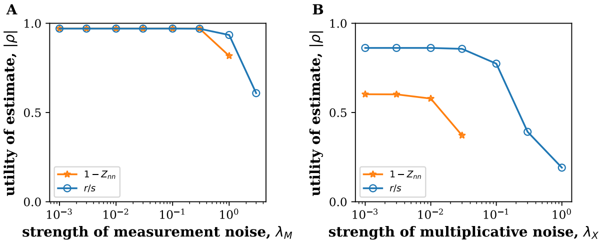

Figure S4: **Estimating search efficacy from noisy data using landscape ruggedness.** Magnitude of the correlation between ruggedness estimated from noisy data and search efficacy remains high even when experimental noise is as large as the community function itself. Two forms of noise were simulated, additive noise in measurement **(A)** and multiplicative noise **(B)**. The strength of the noise $\lambda_M$ and $\lambda_X$ is measured relative to the community function; therefore a noise strength of one means that the contribution from noise is as large as the community function itself. The community function was the productivity of a pair of species in panel A and diversity in panel B. Noise was simulated as described in SI. Data was obtained from simulations in Fig 5.

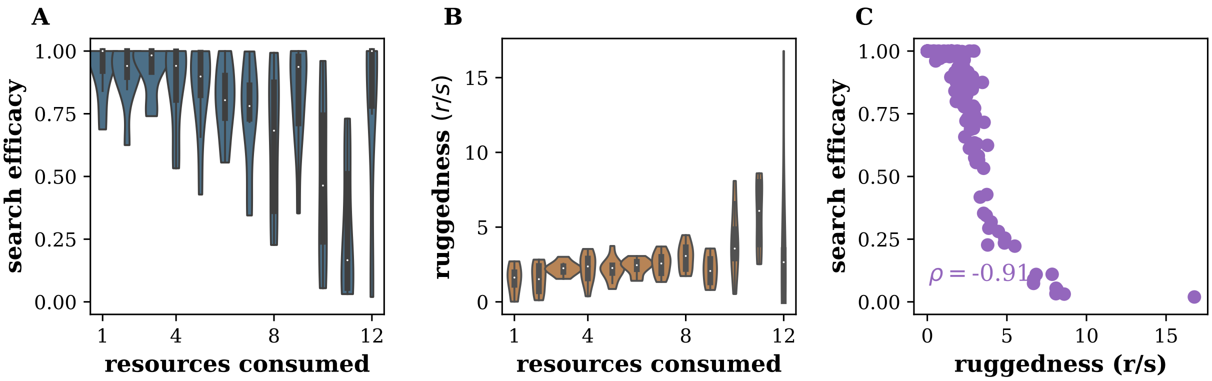

Figure S5: **Niche overlap increases ruggedness and difficulty of search in the presence of cross-feeding. (A,B)** Search outcome and ruggedness of a cross-feeding model where the niche overlap between species was varied by changing the number of resources consumed by each species, mirroring results obtained in model without cross-feeding (Figs. 3, 5). **(C)** Ruggedness remained informative of search efficacy. Community function was the abundance of a focal species. Parameters are described in Methods.

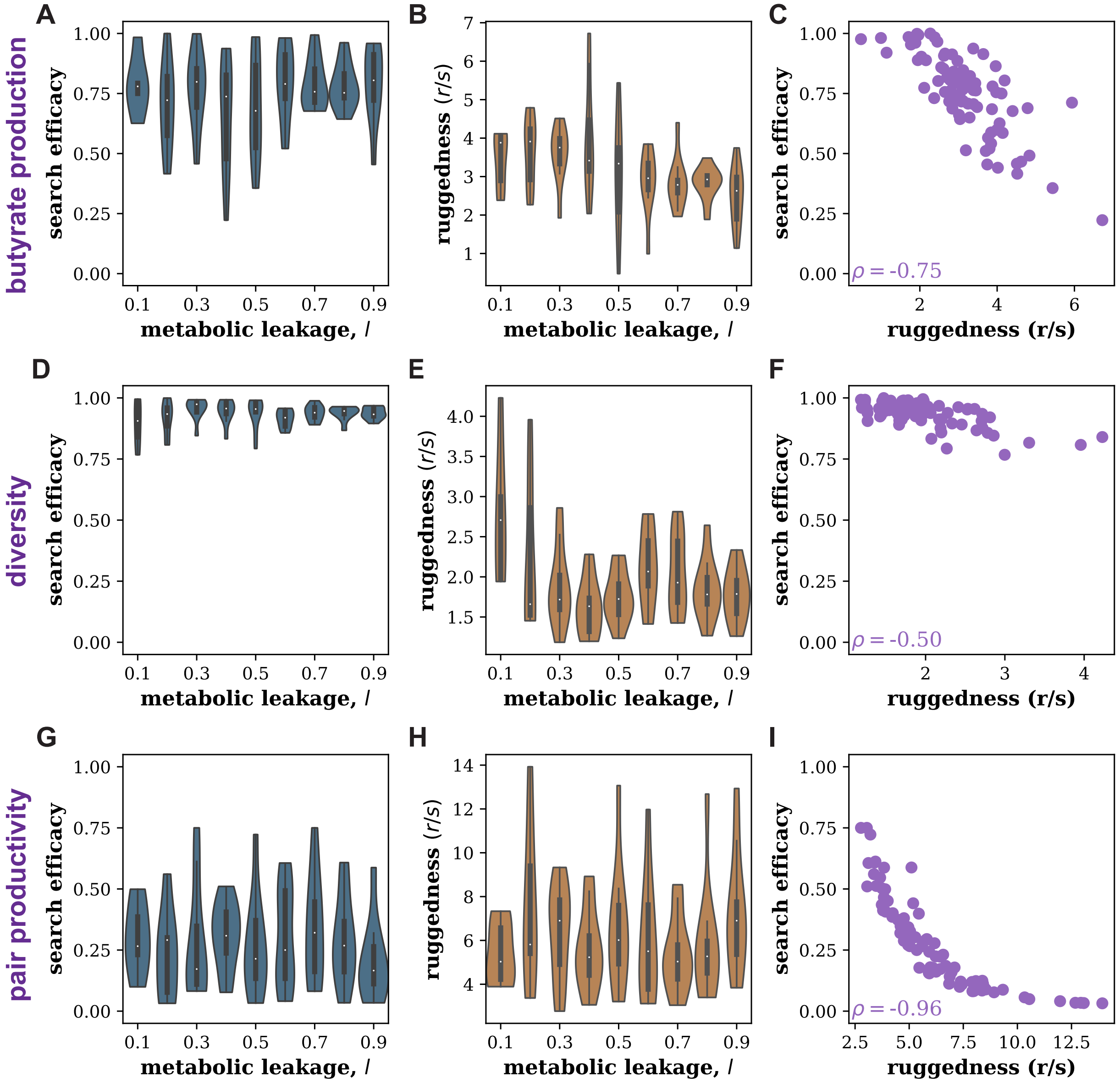

Figure S6: **Ruggedness and search efficacy in the crossfeeding model.** Panels demonstrate the search efficacy, ruggedness, and correlation between search efficacy and ruggedness on landscapes with community functions of butyrate production, diversity, and pair productivity. All reported correlations were statistically significant.

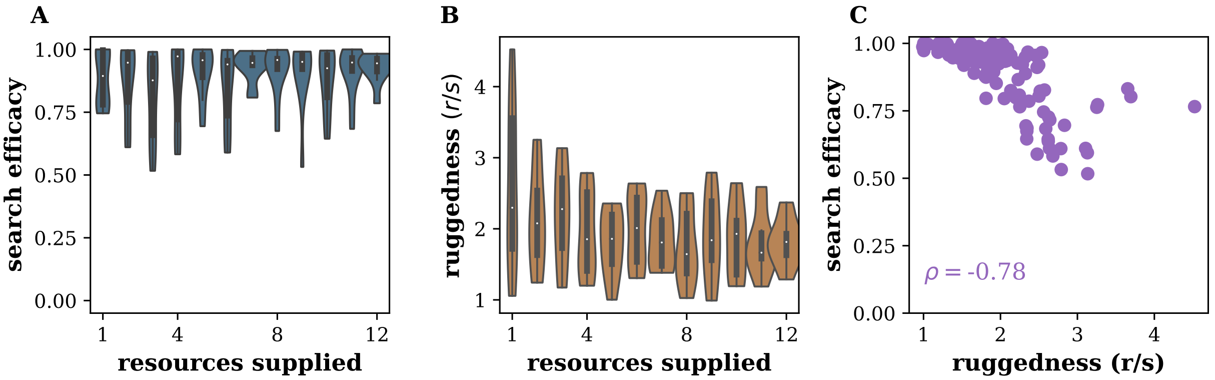

Figure S7: **Landscape ruggedness is informative of search efficacy at different levels of environmental complexity. (A,B)** Search efficacy and ruggedness of a cross-feeding model where the number of resources supplied to the community was varied. **(C)** Ruggedness remained informative of search efficacy. Community function was the abundance of a focal species. The total amount of resources supplied was held fixed in these simulations with parameters as described in Methods. Simulations where the amount of each resource supplied was held fixed instead gave similar results.

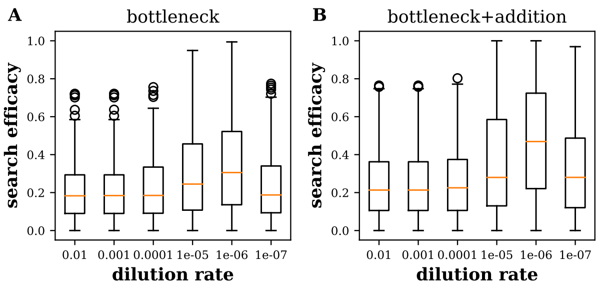

Figure S8: **Performance of dilution-based search protocols is optimal only for a small range of dilution factors.** The dilution-based search protocols, 'bottleneck' and 'addition + bottleneck', have a high search efficacy only if the bottlenecking step kills a few species but not too many. Therefore, it works best for a narrow range of dilution factors where only order 10 cells survive bottlenecking, before being subject to invasion [34]. Community function was the productivity of a pair of species and simulations were the same as in Fig 9

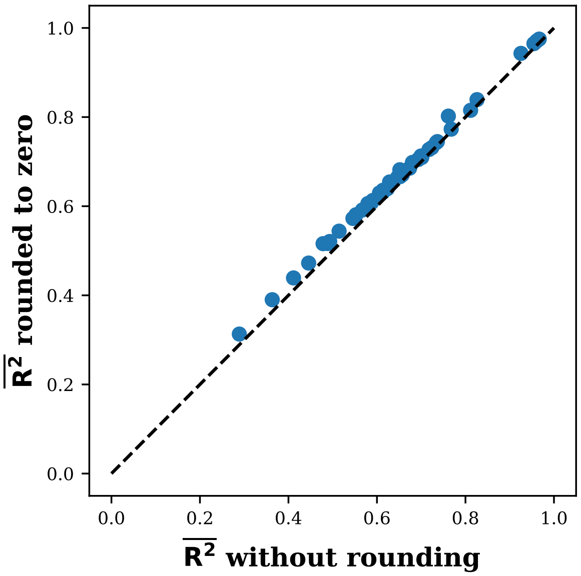

Figure S9: $\overline{R^2}$**, is unaffected by rounding negative predictions of the linear model to zero.** $\overline{R^2}$ after rounding negative abundance predictions to zero is in good agreement with $\overline{R^2}$ computed without rounding negative predictions of the linear model shown in Fig 2.

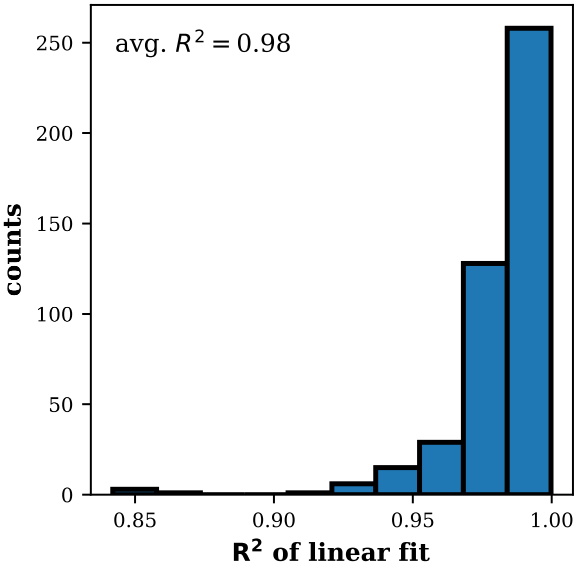

Figure S10: **Rate of metabolic activity is constant over the duration of the experiment.** While the experiment by Langenheder et. al. [35] assayed the cumulative metabolic activity of the microbial communities at different timepoints, used the rate of metabolic activity as the community function. The rate of metabolic activity measured as the slope of a linear fit to the cumulative metabolic activity. The rate was at steady state as evidenced by the high $R^2$ of linear fit to the data as shown in the figure.

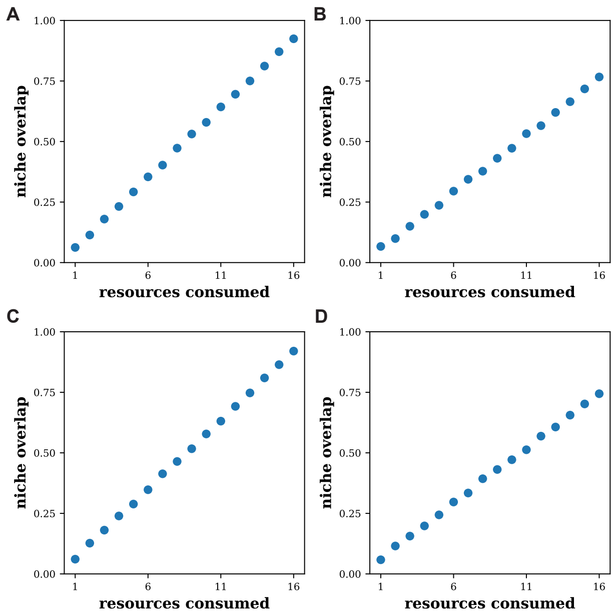

Figure S11: **Niche overlap between species increases with number of resources consumed for different distributions of consumption matrix elements.** The niche overlap in 16 species pools where the non-zero consumption matrix elements were sampled from a uniform distributions **(A,B)**, and lognormal distributions **(C,D)**. Niche overlap was quantified by the average cosine similarity between the consumption vectors of each species pair in the pool, as in previous studies [33]. The uniform distribution in A extended from 0.5 to 1.5; in B extended from 0.1 to 20.5. The lognormal distribution parameters were $\mu = 0$, $\sigma = 0.3$ in panel C and $\mu = 0.6$, $\sigma = 0.6$ in panel D. The extent of increase in niche overlap reduces as the variability in the sampling decreases.

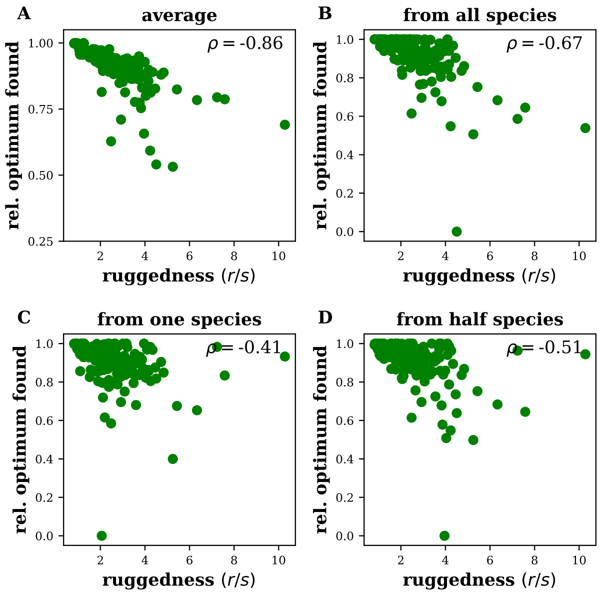

Figure S12: **Ruggedness remains informative of search efficacy even if initial community was fixed.** For Shannon diversity, the ruggedness measure is informative not only about the average search outcome **(A)**, but also about search outcome starting from the community with all species present **(B)**, a randomly chosen species in monoculture **(C)**, and from a randomly chosen community with half the candidate species present **(D)**. Simulation data was the same as in Fig 3.
